# The versatility of Ascorbate Peroxidase-aided mapping uncovers insights of the nuclear lamina interactions and function

**DOI:** 10.1101/2020.02.05.935635

**Authors:** Joseph R. Tran, Danielle I. Paulson, James J. Moresco, Stephen A. Adam, John R. Yates, Robert D. Goldman, Yixian Zheng

## Abstract

The nuclear lamina (NL) is a proteinaceous network found beneath the inner nuclear membrane. The NL is linked to a number of dynamic cellular activities including chromatin organization, transcription and RNA/protein trafficking through nuclear pores. Our understanding of the NL has been hindered in part by the general insolubility and low extractability of proteins from this region. This has spurred the development of proximity ligation methods that label proteins and/or DNA near the NL for systematic identification (Bar et al., 2018; Chen et al., 2018b; Guelen et al., 2008; Roux et al., 2012). To simplify labeling and improve temporal resolution, we fused APEX2 (Hung et al., 2014; Lam et al., 2015) to the nuclear lamina protein lamin-B1 to map proteins, RNA and DNA associated with the NL. We show that APEX2 labeling of the NL is robust and requires as little as 20 seconds. In addition to identifying the NL proteome, this method revealed NL-proximal RNA species that were largely spliced. These NL-proximal RNAs show a bias toward long 3’ UTRs, suggesting an RNA-regulatory role of the NL. This is further supported by the finding of a bias toward longer 3’ UTRs in genes deregulated in lamin-null cells. Interestingly, these RNAs share a sequence motif in their 3’ UTRs. Finally, we demonstrate that the APEX2 method can reliably map lamina-associated domains (LADs) at different stages of the cell cycle, revealing a variability of short LADs regions enriched for histone lysine 27 trimethylation (H3K27me3). Thus the APEX2 method report here is a useful addition to the molecular toolbox for the study of the NL and permits the identification of proteome, transcriptome, and genome elements associated with this nuclear substructure.

## Introduction

The nuclear lamina (NL) is a substructure of the nucleus that resides beneath the inner nuclear membrane. The major structural components of the NL are the intermediate filament proteins, the A- and B-type lamins (Burke and Stewart, 2013; Dechat et al., 2008). The NL may influence the trafficking of RNA and proteins through a lamin meshwork by ensuring an even distribution of NPCs throughout the nuclear envelope (Guo et al., 2014). The NL also organizes the genome by interacting with regions of DNA known as lamina-associated domains (LADs) (Guelen et al., 2008). The study of the NL has gained increased interest due to the distinct human pathologies that are caused by mutant versions of NL proteins and the connections of lamins to aging (Chen et al., 2014; Chen et al., 2015; Hatch and Hetzer, 2014; Schreiber and Kennedy, 2013; Tran et al., 2016; Yue et al., 2019). However, the importance of the NL in general organismal biology was first highlighted by the discovery of differential expression of A- and B-type lamins during embryonic development (Rober et al., 1989; Stewart and Burke, 1987) and further punctuated by the discovery of the essential nature of NL genes for viability and overall genome organization (Coffinier et al., 2010; Kim et al., 2011; Sullivan et al., 1999; Vergnes et al., 2004). More recently, the NL has been implicated in aging with the apparent reduction or alteration of lamin proteins in aged animals and in a form of cellular aging termed senescence (Chen et al., 2014; Chen et al., 2015; Freund et al., 2012; Frost et al., 2016; Lattanzi et al., 2014; Shimi et al., 2011; Tran et al., 2016; Yue et al., 2019). Studies of the NL in these contexts have revealed functions for the NL in highly dynamic processes such as genome organization, gene transcription, signal transduction, protein/RNA trafficking, and cell division. However, our understanding of these NL functions is rather limited, in part due to an incomplete characterization of the molecular components of the NL and our limited knowledge of the dynamics associated with it.

The study of the NL is challenging as it has long been recognized as a proteinaceous structure with limited solubility (Gerace et al., 1984; Moir et al., 2000). This insolubility occurs throughout most of a cell’s life with the exception of animal cell mitosis when the nuclear envelope is disassembled. Early biochemical studies of the NL employed fractionation of this structure, but this method is limited by the ability to screen for different biological molecules and the amount of starting material required. This problem led to the development/utilization of a number of proximity ligation methods such as Biotin-mediated identification (BioID), DNA adenine methylation-mediated identification (DamID), Tyramide Signal Amplification (TSA) and Biotinylation by antibody recognition (BAR) (Bar et al., 2018; Chen et al., 2018b; Guelen et al., 2008; Roux et al., 2012). The principle behind each of these methods is the enzymatic tagging of proteins and/or nucleic acids in the proximity of the NL with a molecule that is either readily identified or amenable to purification. As examples, DamID uses a DNA adenine methyltransferase to label DNA at the NL and BioID utilizes a promiscuous BirA biotin ligase to label NL proteins with biotin (Guelen et al., 2008; Roux et al., 2012). More recently the TSA method, classically used to increase the signal of immunostaining procedures by regional horseradish peroxidase-catalyzed deposition of biotin, was used to map the NL-associated proteome (BAR method) and lamina-associated domains (LADs, TSAseq) (Bar et al., 2018; Chen et al., 2018b). These proximity methods, while extremely useful, do have some restrictions. DamID for example, targets DNA and does not provide proteomic information (van Steensel and Henikoff, 2000) while TSA/BAR requires specific antibodies which might not discriminate between isoforms or genetic variants (Bar et al., 2018; Chen et al., 2018b). Further, some of these approaches have limited temporal resolution, or they require large amounts of cellular starting material. An alternative enzyme suitable for the study of the NL, called Ascorbate Peroxidase (APEX), was recently developed by the Ting group for the purposes of proteomic and RNA identification (Fazal et al., 2019; Hung et al., 2014; Kaewsapsak et al., 2017). This enzyme, which has been extensively engineered into the highly-reactive form, APEX2 (Lam et al., 2015), uses hydrogen peroxide to catalyze the covalent addition of a radicalized biotin-phenol moiety to both protein and RNA species. The APEX2 enzyme is reported to have a labeling radius of around 20 nm, making it suitable for the study of structurally-defined regions such as the NL (Martell et al., 2012).

Here, we describe the use of APEX2 to obtain a more complete picture of the NL by the identification of its associated proteins, RNA and DNA. We show that the NL associates with proteins involved in RNA regulation such as mRNA splicing and stability, and that the APEX identified NL proteome exhibits strong overlap with that identified by the related BAR method (Bar et al., 2018). The use of APEX to identify NL-associated RNA species suggests an interesting role for the NL in the regulation of a select group of mRNAs. Finally, the APEX2 method allows easy study of LADs in different cell cycle stages.

## Material and Methods

### Cell culture

HEK293FT (Thermo, #R70007), HCT116 (ATCC, #CCL-247) and K562 (ATCC, #CCL-243) cell lines were cultured in DMEM, McCoy’s 5a and RPMI media, respectively. The media was supplemented with 10% FBS for DMEM and McCoy’s 5a and 15% FBS for RPMI. The cells were grown at 37 °C in 5% CO2.

### Plasmid construction and transfection

The APEX2-lamin-B1 fusion was generated by replacing the tubulin cDNA in Addgene plasmid #66171 (Lam et al., 2015) with the human lamin-B1 cDNA using the XhoI and BamHI cloning sites. A (GGGGS)3 linker was placed between the APEX2 and lamin-B1 (Huston et al., 1988). Transfections were performed using Lipofectamine 2000 (Thermo, #11668030) according to the manufacturer’s instructions. The APEX2-lamin-B1 construct was expressed in cells for 24-48h. The plasmid is available on Addgene #139442.

### FACS sorting

Cells were harvested by trypsinization and neutralized in medium containing FBS. The cells were fixed for 10 minutes at room temperature by adding freshly prepared 4% PFA to a final of 1%. The fixation was quenched with 125 mM glycine and the cells were washed with PBS. Cells were resuspended in phenol-red free HBSS + 2% FBS and stained with Hoechst 33342 for at least 20 minutes at room temperature before FACS sorting for G1, S, G2/M phases of the cell cycle.

### APEX2 reaction

The APEX2 reaction was done under either previously published unfixed (standard) (Hung et al., 2014) or PFA-fixed conditions, depending on the material to be isolated. For protein identification by mass spectrometry, cells expressing the APEX2-lamin-B1 construct were incubated with 6 ml of media 250 μM biotin-phenol (Iris Biotech, #LS-3500.0250) for 30 minutes at 37 °C. An equal volume of 2 mM H_2_O_2_ was added and incubated for 1 minute at room temperature. The reaction was quenched by adding a solution containing Trolox (Sigma, #238813) and sodium ascorbate (Sigma #PHR1279) to a final of 5 mM and 10 mM, respectively. The solution was aspirated and the cells were immediately lysed on the dish with 1 ml of RIPA buffer (50 mM Tris, 150 mM NaCl, 0.1% (wt/vol) SDS, 0.5% (wt/vol) sodium deoxycholate and 1% (vol/vol) Triton X-100, pH 7.5) containing a protease inhibitor cocktail tablet (Sigma cOmplete, EDTA-free, #11873580001). For isolation of RNA-bound biotinylated proteins under unfixed conditions, cells were treated as above except lysis was done in the presence of 100 units/ml RNasin (Promega, #N251B).

For RNA and DNA pull downs using the RNA- and DNA-bound biotinylated proteins, cells were harvested by trypsinization and neutralized with medium containing 10% FBS. The cells were then pelleted and fixed for 10 minutes at room temperature by adding freshly prepared 4% PFA to a final of 1%. The fixation was quenched with 125 mM glycine and the cells were washed with phosphate buffered saline (PBS), pH 7.2. The cells were then permeabilized with PBS + 0.25% Triton X-100 for 5 minutes at room temperature. In the case of RNA, all solutions were supplemented with 100 units/ml RNasin. The cells were briefly pelleted and washed with PBS. The pellet was resuspended in 100 μl of DMEM medium containing 10% FBS, due to the requirement for heme in the reaction (Martell et al., 2012), and 100 μM biotin-phenol for 5 minutes at room temperature. An equal volume of 2 mM H_2_O_2_ (final concentration 1mM) was added to the solution, mixed and incubated for 20 seconds. The reaction was then quenched with Trolox (5 mM) and sodium ascorbate (10 mM). A small aliquot (40 μl) of the APEX reaction was stained and examined by fluorescence microscopy (see below) to confirm the reaction. The remainder of the APEX reaction could be used immediately for isolation of material or snap frozen and stored at −80 °C for later use.

### Fluorescence microscopy

For initial testing of the APEX reaction, cells were grown on coverslips and transfected with the APEX2-lamin-B1 construct. The reaction was performed either with live cells, or after 1% PFA fixation as described above. For examination of diffusion of biotinylated material in living cells, the APEX reaction was first performed and the cells were washed twice with media containing Trolox and sodium ascorbate, twice with standard growth medium and then cultured for the indicated times in standard growth medium. The cells were then fixed and stained. The APEX reaction was confirmed by incubating with mouse-anti FLAG-M2 antibody (1:1000, Sigma, #F3165) overnight at 4 °C and followed by incubation with Streptavidin conjugated to Alexa 488 (1:200, Biolegend, #405235) and an anti-mouse secondary conjugated (1:1000) to an Alexa fluorophore. Other primary antibodies used for immunofluorescence include rabbit anti-HNRNPA1 (1:2500, ProteinTech, #11176-1-AP), rabbit anti-SRSF1 (1:2500, ProteinTech, #12929-2-AP), rabbit anti-SRSF7 (1:2500, Novus, #NBP1-92382). For cell suspensions, all washing procedures were done by pelleting at 500g x 5m and resuspended in Prolong anti-fade gold for mounting and imaging on an SP5 confocal microscope (Leica).

### Isolation of protein, RNA, and DNA

The isolation of protein, RNA, and DNA was performed using the same precipitation pipeline with only minor differences that depended on the desired molecules. For all protocols, cells were initially lysed in RIPA buffer containing a protease inhibitor cocktail tablet (Sigma cOmplete, EDTA-free) for 30 minutes with rotation at 4 °C. In the specific case of RNA isolation, the RIPA buffer was also supplemented with RNasin (100 units/ml). In the case of DNA isolation, the resulting lysate was sonicated using a Diagenode Bioruptor Pico, cleared by centrifugation at 15,000g x 30 seconds, and at this point 1/10^th^ of the lysate was obtained for the input fraction. Lysates, regardless of target were incubated with 70 μl streptavidin magnetic beads (Pierce, # 88817) overnight at 4 °C using end-over-end rotation. The next day, the magnetic beads were harvested on a magnetic rack and washed in sequence with 2 x 1 ml RIPA, 1 x 1 ml with high salt buffer (1M KCl, 50mM Tris-Cl pH 8.0, 5mM EDTA), 1 x 1ml Urea wash buffer (2M Urea, 10mM Tris-Cl pH 8.0), and then 1 x 1ml RIPA (Hung et al., 2014). For RNA experiments, the streptavidin beads were treated with a solution of 0.1M NaOH and 0.05M NaCl to remove RNases and cleared with a solution containing 0.1 NaCl prior to use and the Urea buffer wash was excluded. The material was subjected to Streptavidin beads-mediated Pull Down (StrePD) followed by processing for each specific target. For protein, the StePD beads were resuspended in 1x SDS-PAGE sample buffer and subject to western blotting or mass spectrometry (see below). For RNA, RNase-free DNase I (Sigma, #716728001) was added to the StrePD beads and incubated at 37 °C for 10 minutes followed by an incubation with Proteinase K at 60 °C for 30 minutes. The RNA was then extracted using Trizol reagent according to the manufacturer’s protocol. For DNA, RNase A (Qiagen, #19101) was added to the StrePD beads and incubated at 37 °C for 10 minutes followed by an incubation with Proteinase K (Takara, #9034) at 60 °C overnight. Both StrePD pulldown DNA and DNA from the input lysate were extracted using Ampure XP beads (Agencourt, #A63881). Quantitation of nucleic acids was done using the Qubit system (Thermo).

### Nuclear and cytosol preparation for RNA-seq

Plasma membranes were disrupted by gently resuspending cell pellets in 10mM HEPES pH 7.5, 60mM KCl, 1mM EDTA pH 8.0, 1mM DTT, 1mM PMSF, 0.075% v/v IGEPAL CA-630 supplemented with 100 units/ml RNasin and rotating the mixture at 4 °C for 5 minutes. The volume of lysis buffer used was approximately 5 times the cell pellet volume. The nuclei were pelleted at 200g for 5 minutes. Half of the cytosolic supernatant was carefully removed from the top and pelleted a second time as initially done and the resulting supernatant was used as the cytosolic fraction. To obtain nuclei, the remainder of the cytosolic fraction was removed and the pellet was washed twice with 10mM HEPES pH 7.5, 60mM KCl, 1mM EDTA pH 8.0, 1mM DTT, 1mM PMSF supplemented with an 100 units/ml RNAsin. The nuclei were then resuspended in the starting lysis volume with lysis buffer. One half of the nuclear fraction was used for RNA extraction using Trizol reagent, followed by quantification by Nanodrop (Thermo).

### Western blotting and mass spectrometry

The APEX reaction was performed as described above and lysed in RIPA. Aliquots were taken for input, post-Streptavidin pulldown and Streptavidin pulldown. Proteins were separated on an SDS-PAGE gel and transferred to nitrocellulose for western blotting with the indicated reagents/antibodies. Detection reagents used were Streptavidin-HRP (1:1000, GE Healthcare, # RPN1231V), rabbit anti-lamin-B1 (1:10000, Abcam, #ab16048), mouse anti-lamin-A/C (1:5000, Active Motif, #39287), rabbit anti-Emerin (1:5000, Santa Cruz, #sc-15378), mouse anti-SC-35 (1:2500, Sigma, #S4045), rabbit anti-HNRNPA1 (1:2500, ProteinTech, #11176-1-AP), rabbit anti-SRSF1 (1:2500, ProteinTech, #12929-2-AP).

### Mass Spectrometry

Proteins were precipitated with 23% TCA and washed with acetone. Protein pellets were solubilized in 8 M urea, 100 mM Tris pH 8.5, reduced with 5 mM Tris(2-carboxyethyl)phosphine hydrochloride (Sigma-Aldrich), and alkylated with 55 mM 2-Chloroacetamide (Fluka Analytical). Digested proteins were analyzed by four–step MudPIT using an Agilent 1200 G1311 quaternary pump and a Thermo LTQ Orbitrap Velos using an in-house built electrospray stage (Wolters et al., 2001).

Protein and peptide identification and protein quantitation were done with Integrated Proteomics Pipeline - IP2 (Integrated Proteomics Applications). Tandem mass spectra were extracted from raw files using RawConverter (He et al., 2015) with a monoisotopic peak option. Peptide matching was done against a reviewed Uniprot human protein database (released 1/22/2014, 20275 entried) with common contaminants and with reversed sequences using ProLuCID (Peng et al., 2003; Xu et al., 2015) with a fixed modification of 57.02146 on cysteine and differential modification of 363.146012 on tyrosine. Peptide candidates were filtered using DTASelect, with these parameters; -p 1 -y 1 --trypstat --pfp 0.01 -DM 10 --DB --dm -in -t 1 (Tabb et al., 2002). GO-term analysis was done using Panther (Mi et al., 2019).

### Cell cycle H3K9me3 Chromatin Immunoprecipitation-sequencing (ChIP-seq)

Asynchronously growing HCT116 cells were fixed with 1% PFA and FACS sorted to obtain ∼1 million cells for each of the cell cycle stages G1, S and G2/M. The cells were lysed with RIPA buffer (see above) supplemented with 500 μM PMSF for 30 minutes with rotation at 4 °C. The lysate was sonicated using a Diagenode Bioruptor Pico and immunoprecipitated with anti-H3K9me3 (Abcam, ab8898) complexed to Protein A/G dynabeads. The precipitates were washed with a low-salt buffer (20mM Tris-Cl, pH 8.0, 150mM NaCl, 2mM EDTA, 1% Triton X-100, 0.1% SDS), high-salt buffer (20mM Tris-Cl, pH 8.0, 500mM NaCl, 2mM EDTA, 1% Triton X-100, 0.1% SDS) and a LiCl buffer (10mM Tris-Cl, pH 8.0, 250mM LiCl, 1mM EDTA, 1% deoxycholic acid, 1% IGEPAL-CA-630) supplied from the Millipore Chromatin Immunoprecipitation kit (Cat #17-295). Precipitates were resuspended in RIPA buffer and digested with Proteinase K overnight at 60 °C. DNA for both input and ChIP was recovered using Ampure XP beads.

### RNA and DNA sequencing

RNA library building was done using the Illumina TruSeq RNA library kit v2 (Illumina #RS-122-2201) with Ribo-depletion. DNA libraries were prepared using the Rubicon Genomics ThruPlex kit (Rubicon Genomics, #R400428). Sequencing was performed on the Illumina NextSeq 500 platform (Illumina). Raw data will be made available through the NCBI Gene Expression Omnibus.

### Data analysis

RNA-seq data was aligned using Bowtie 2.26 and Tophat2 with default settings using the hg19 assembly. Counting into features was done using featureCounts from the Subread package v1.5.2 (Liao et al., 2014) using the -s 0 parameter. Exonic and intronc read counts were done using RefFlat coding exons, and RefFlat introns. We removed any intron containing snoRNAs, miRNA and/or lincRNAs. Wild-type and lamin-null RNAseq data was obtained from GEO GSE89520 (Zheng et al., 2018). Differential enrichment was calculated using the glmFIt method in edgeR v3.42.3 R package (Robinson et al., 2010). We used a threshold of an FDR less than or equal to 0.05 to determine fraction enrichment. Plotting was done in either Rstudio v0.98.953 (R v3.5.1) using pHeatmap v1.0.12 or Microsoft Excel 2016. Motif analysis was done using MEME v5.1.0 with -mod zoops (for RIP), -mod anr (for mESC data), -minw 5, -maxw 10 and - markov_order 0 settings (Bailey and Elkan, 1994). Browser tracks were displayed in Integrative Genomics Viewer (IGV) v2.3.94.

LADs mapping was done by first aligning input and StrePD reads using the hg19 assembly and Bowtie 2.26 and then called into 100 kilobase genomic windows using the coverage function in Bedtools v2.26.0. Enrichment was calculated by first normalizing the StrePD and input data by read count to one million and then transforming the StrePD/input ratio by log2. LADs (DamID and TSAseq) data from K562 was obtained from GEO Omnibus GSE66019 (Chen et al., 2018b). K562 RNA-seq data was obtained from GEO Omnibus GSM958731 and epigenome data for HCT116 cells was obtained from the ENCODE project (HCT116 reference epigenome series ENCSR361KMF). For H3K9me3 ChIP data, input and ChIP reads were aligned using Bowtie 2.26 and the hg19 assembly. Peak calling was performed using MACS2 v2.1.1.20160309 (Zhang et al., 2008) using default settings. Quantitation of histone signal in LADs was done using the bigwigAverageOverBed function in kentutils v3.62. Correlative analysis, Hidden Markov Model (HMM), statistics and graphical plotting were done either in R Studio using the pHeatmap v1.0.12, corrplot v0.84, Hmisc v4.2-0, and depmixs4 v1.4-0 packages or Microsoft Excel 2016. Browser tracks were displayed using the UCSC Genome Browser with a smoothing window of 2-3.

## Results

### APEX2-lamin-B1 labels the nuclear periphery

We transfected HEK293FT cells using no plasmid or plasmid expressing FLAG-APEX2-lamin-B1 (human lamin-B1) and performed the APEX2 labeling reaction using the protocol similar to that described by Ting and colleagues (Figure 1A Method #1) (Hung et al., 2016). The cells were incubated with biotin-phenol in their culture medium and then treated with hydrogen peroxide for 1 minute prior to processing. After cell fixation, we performed Streptavidin labeling and immunostaining using the Flag-M2 tag. We found that the APEX2 reaction is very robust, but the streptavidin signal is diffuse throughout the entire nucleus, whereas the Flag-M2 tag staining showed a clear nuclear lamina localization (Figure 1B). We tried limiting diffusion by reducing the APEX2 reaction down to 15 seconds, but continued to observe a similar diffuse-staining pattern for streptavidin (Figure S1A). This suggests that APEX2 on lamin-B1 might have labeled nearby nuclear proteins that can diffuse throughout the nucleus in live cells prior to fixation. We next determined whether the APEX2 reaction can be performed on fixed cells (Figure 1A Method #2). We found fixing cells with 1% paraformaldehyde (PFA) prior to permeabilizing with Triton X-100 followed by incubation with biotin-phenol and hydrogen peroxide resulted in a strong streptavidin staining that coincided with FLAG-M2 staining for the APEX2-lamin-B1 fusion protein (Figure 1C, white arrows). This result demonstrates that APEX2-based biotinylation can be performed in fixed cells, which captures both stable and transient NL-associated proteins.

**Figure 1:**
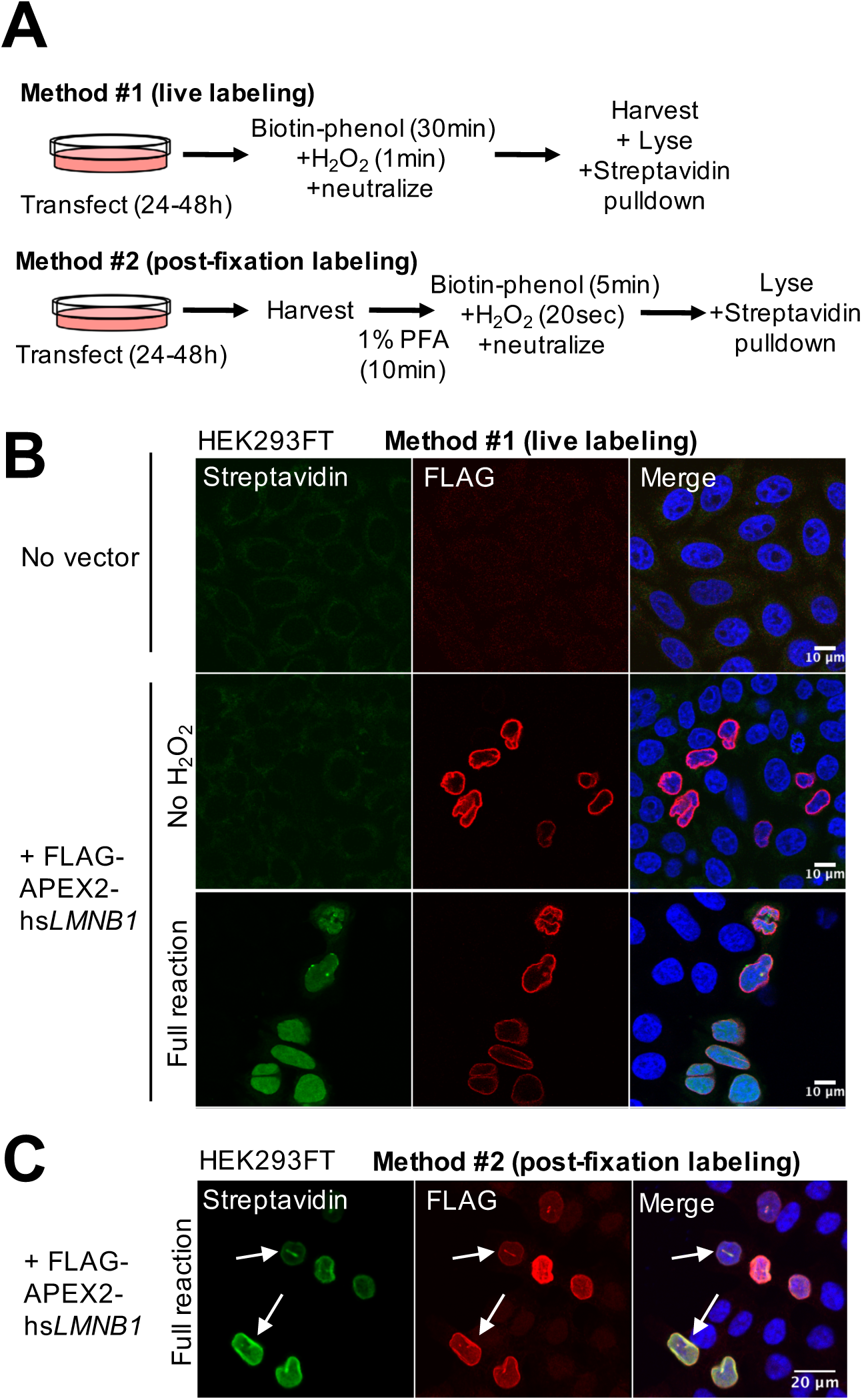
APEX2-lamin-B1 labels the nuclear lamina. (A) The experimental pipeline for live and paraformaldehyde (PFA)-fixed cells used for APEX2 reactions in this study. Labeling time is indicated in parentheses. (B-C) Fluorescence streptavidin staining of biotinylated proteins/RNA by APEX2 labeling in live HEK293FT cells (B) and PFA fixed HEK293FT cells (C). DAPI (DNA), Streptavidin (biotinylated material) and FLAG M2 (marking the Flag-tagged APEX2-lamin-B1). Arrows point toward subtle variations in both Streptavidin and FLAG-M2 staining.

The robustness of the APEX2 enzyme under fixation conditions prompted us to examine the stability of the enzyme under other conditions. We found that the APEX2 enzyme was still reactive in fixed cells that were stored at 4 °C for up to three weeks (Figure S1B) and that the enzyme also withstood flash freezing in liquid N_2_ (Figure S1C). Finally, we performed the APEX-reaction in live cells, neutralized the reaction and chased the biotinylated proteins for up to an hour. We found that the biotinylated nuclear protein did not exit the nucleus (Figure S1D). Our results show that the APEX2 reaction is robust under different experimental and cell fixation conditions permitting the biotinylation of NL-associated nuclear proteins; and in unfixed cells the diffuse nuclear signal was likely due to diffusion of biotinylated protein inside the nucleus.

### APEX2-lamin-B1 labeling identifies NL-associated RNA with long 3’ UTR

Previous studies have used the APEX procedure to isolate RNAs (APEX-RIP) associated with cellular organelles and structures. This APEX-RIP procedure can be done by precipitating protein/RNA complexes, or by directly precipitating RNA since the APEX reaction will label RNA (Fazal et al., 2019; Kaewsapsak et al., 2017). In our experiments, we chose to precipitate protein/RNA complexes in order to isolate NL-enriched RNAs since this is expected to yield a more extensive dataset. We generated RNA sequencing (RNA-seq) datasets for total nuclear and total cytosol RNA, and then used native APEX-lamin-B1-RIP and PFA-fixed APEX-lamin-B1-RIP to obtain NL-associated RNA (Figure 2A, Supplementary Table 1). We included cell fixation followed by APEX2 labeling because it captures RNA-protein complexes when they are restricted at the NL, thus adding confidence that the RNAs identified are indeed NL associated. The biological replicates for each experimental condition are consistent with Pearson correlation values ranging from 0.84 to 1.00 (Figure S2A, S2B). We first compared the nuclear and cytosolic RNA-seq fractions (genes with >5 counts per million) and found the expected enrichment (∼15-20-fold over cytosolic reads) of known nuclear residents, *XIST*, *MALAT1* and *NEAT* (Figure S2C, left panel) (Hutchinson et al., 2007). We found a total of 3596 cytosolic- and 5073 nuclear-enriched RNAs using differential expression analysis with an FDR cutoff of 0.05 from two replicate experiments (Figure S2C, right panel).

**Figure 2:**
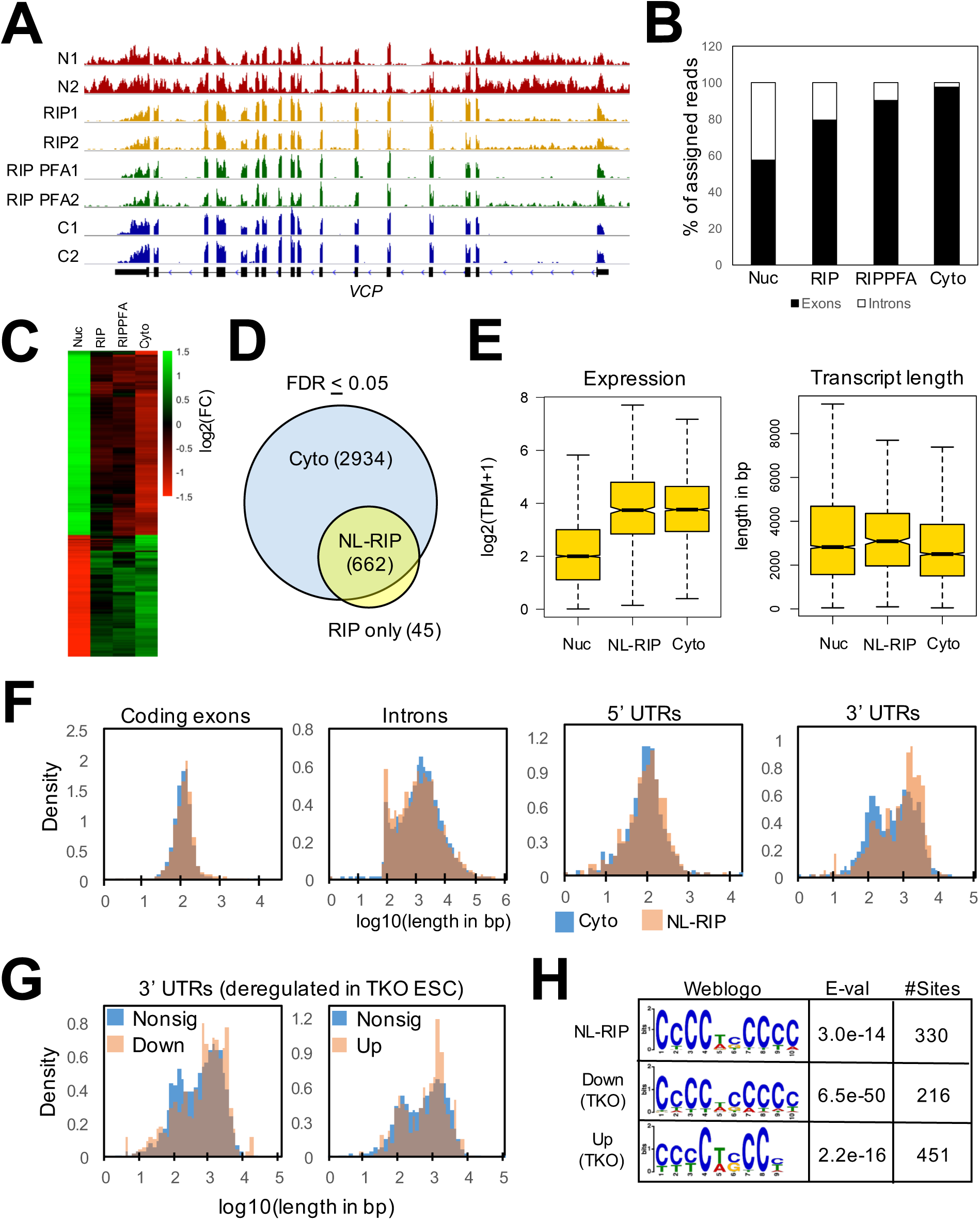
APEX2-lamin-B1 identifies RNA-species associated with the NL. (A) Integrative Genomics Viewer (IGV) browser tracks of RNA-seq for two biological replicates of nuclear RNA “N1, N2”, RNA by RIP of live cells “RIP1, RIP2”, RNA by RIP of PFA-fixed cells “PFA-RIP1, PFA-RIP2” and cytosol RNA “C1, C2”. The *VCP* locus (hg19) is shown. These data were generated from the HEK293FT cell line. (B) Average exon and intron content for each dataset presented as a percentage of reads mapped to either feature. (C) Heatmap display of the log2 fold change for genes enriched in each fraction with an FDR of less than or equal to 0.05 compared to the nucleus. Nuclear enrichment was calculated against the cytosol (D) Venn diagram showing the overlap between the RIP enriched dataset and the cytosol-enriched datasets. Overlap between the cytosolic and RIP dataset is labeled as “NL-RIP” while Cytosolic and RIP only enrichment is labeled as “Cyto” and “RIP only”, respectively. For subsequent analysis, we excluded the RIP only group because of the low number of mRNAs. (E) Boxplots showing the expression level (left panel) of genes and gene length (right panel) for RNAs enriched in each fraction. The notches represent the 95% confidence interval around the median. (F) Density histograms comparing the size distribution of different RNA features from our NL-RIP and Cyto datasets. We examined coding exons, introns, 5’ UTRs and 3’ UTRs as curated by UCSC Genome Browser. (G) Density histograms comparing the distribution of sizes for the 3’ UTR of deregulated genes in the lamin-null (TKO) RNA-seq dataset. (H) MEME motif search showing a similar C-rich motif enriched in the 3’ UTRs of the NL-RIP fraction and in genes deregulated in TKO mESC.

As expected, the nuclear RNA-seq dataset had ∼42% of reads map to introns while only ∼2% of cytosolic reads are intronic (Figure 2B). Interestingly, both native APEX2-lamin-B1-RIP and fixed APEX2-lamin-B1-RIP identified RNAs with low intronic reads (Figure 2A, 2B) and the intron/exon profile is similar to that seen in the cytosolic RNAs. Enrichment analysis of RNAs in APEX-lamin-B1-RIP versus the nuclear RNAs also indicates that RNAs from APEX-RIP are similar to the cytosolic RNAs (Figure 2C). This shows that the NL-associated RNAs are spliced and *en route* to the cytosol. Consistent with this, our mass spectrometry data detect mainly nucleoplasmic nuclear pore components (TPR and NUP153) (see more below and Supplementary Table 2) (Frosst et al., 2002; Sukegawa and Blobel, 1993). Additionally, we observed no detectable diffusion of biotinylated material outside of the nucleus in our live-cell APEX label and chase experiments (Figure S1D), which further support the idea that the APEX labeling captured the NL-associated RNAs. We found that previously identified mRNA transcripts with retained introns in the nuclear and APEX-RIP samples which are then mostly, but not completely removed in the cytosolic RNA-seq fraction (Figure S2D, black bars) (Boutz et al., 2015; Fazal et al., 2019; Lareau et al., 2007).

To characterize RNA species that are enriched at the NL, we intersected significantly different genes (FDR < 0.05) from both of our APEX-RIP and PFA-fixed APEX-RIP datasets to yield a consensus set of 707 NL-enriched RNAs (Figure S2E). These NL-enriched RNAs were protein-coding mRNAs. We did not observe any enrichment for noncoding RNAs (HUGO-defined) although they were present in the RNA-seq data (Supplementary Table 1). Most these NL-enriched mRNAs (93.4%) are found in the cytosolic-enriched population (Figure 2D, “NL-RIP”), but the reverse is not true with only 18.4% of cytosolic-enriched RNAs being NL-enriched mRNAs. We found a small population of mRNAs (45) that were exclusively enriched in the RIP fraction (Figure 2D, “RIP only”) and excluded these from further analysis due to the low number. Next, we compared the expression levels and the overall length of the mRNAs among the NL-RIP, cytosolic (Cyto) and nuclear (Nuc) fractions. We found that mRNAs in the NL-RIP and cytosolic fractions are expressed at higher levels than those of the Nuclear fraction, but mRNA lengths are similar among all three fractions (Figure 2E). When we compare specific features of the mRNAs, we find that the NL-RIP population shows a bias toward longer 3’ UTRs, but other features such as the 5’ UTRs, coding exons and introns were similar (Figures 2F and S2F). This suggests that a subset of mRNAs with long 3’ UTRs associate with the NL before being exported into the cytosol.

Next, we examined the up- and down-regulated genes in our previously published wild-type and lamin-null (triple knockout of lamin-A/C, -B1, and -B2, TKO) mouse embryonic stem cells (mESC) RNAseq datasets (Zheng et al., 2018). We found that 3’ UTRs in both the up- and down-regulated genes were larger than the unchanged RNA population (Figure 2G), but other features were similar in size (Figure S2G). Taken together, this suggests that RNA regulation via 3’ UTRs could explain some of the differentially expressed genes upon lamin deletion in mESCs. In an effort to identify potential regulatory motifs, we performed a MEME search of non-redundant 3’ UTRs from APEX NL-RNAs that were greater than the median size. This search identified an enrichment for a short nucleotide motif high in C residues (CCCCWCCCC, W can be A or U, Figure 2H). We next performed a motif search of down- and up-regulated genes in TKO mESCs compared to the WT mESCs and found a significant enrichment for a similar C-rich motif (Figure 2H). Thus, the APEX-RIP method of identifying NL-associated RNAs reveals a potential role for the NL in interacting and regulating some transcripts with long 3’ UTR that contain C-rich motifs.

### APEX2-lamin-B1-based labeling identifies potential NL-associated proteins with RNA splicing and stability functions

To explore if APEX2-lamin-B1 can be used to identify an NL-associated proteome, we performed the APEX reaction with unfixed cells prior to protein purification to facilitate mass spectrometry. Biotinylated proteins were observed and could be efficiently precipitated by streptavidin beads (“StrePD”) (Figure 3A). As anticipated, known NL proteins (lamin-A/C and Emerin) were detected by western blotting (Figure 3A). Mass spectrometry identified 338 proteins with an average of 5 or more spectral counts from two replicate experiments (Supplementary Table 2). To assess the reliability of our proteomic data, we compared our dataset with a recently published NL proteomic dataset generated by tyramide signal amplification (here referred to as “TSA-BAR”) (Bar et al., 2018) and found 46.7% overlap (Figure S3A). Our dataset also shows a reasonable overlap with many previous datasets (Figure S3A) generated by different methods (Bar et al., 2018; Depreux et al., 2015; Dittmer et al., 2014; Dreger et al., 2001; Engelke et al., 2014; Fu et al., 2015; Kubben et al., 2010; Roux et al., 2012; Schirmer et al., 2003; Thul et al., 2017). We next compared our dataset against a list of 120 proteins identified in at least three previous NL proteomic experiments and found that 66 (55%) of these proteins are in our APEX study (Supplementary Table 1, Figure 3B, black circles). We note that our APEX-based NL mass spec candidates did not contain many secreted proteins (3/338) or transmembrane proteins (27/338, Figure S3B). A select number RNA splicing proteins (e.g., SC-35, SRSF1 and HNRNPA1) found in our mass spectrometry were independently validated by western blot experiments (Figure S3C). Immunostaining showed that a sub-fraction of several splicing factors, HNRNPA1, ASF1 and SFRS7, are localized at the NL (Figure 3C, inset arrowheads).

**Figure 3:**
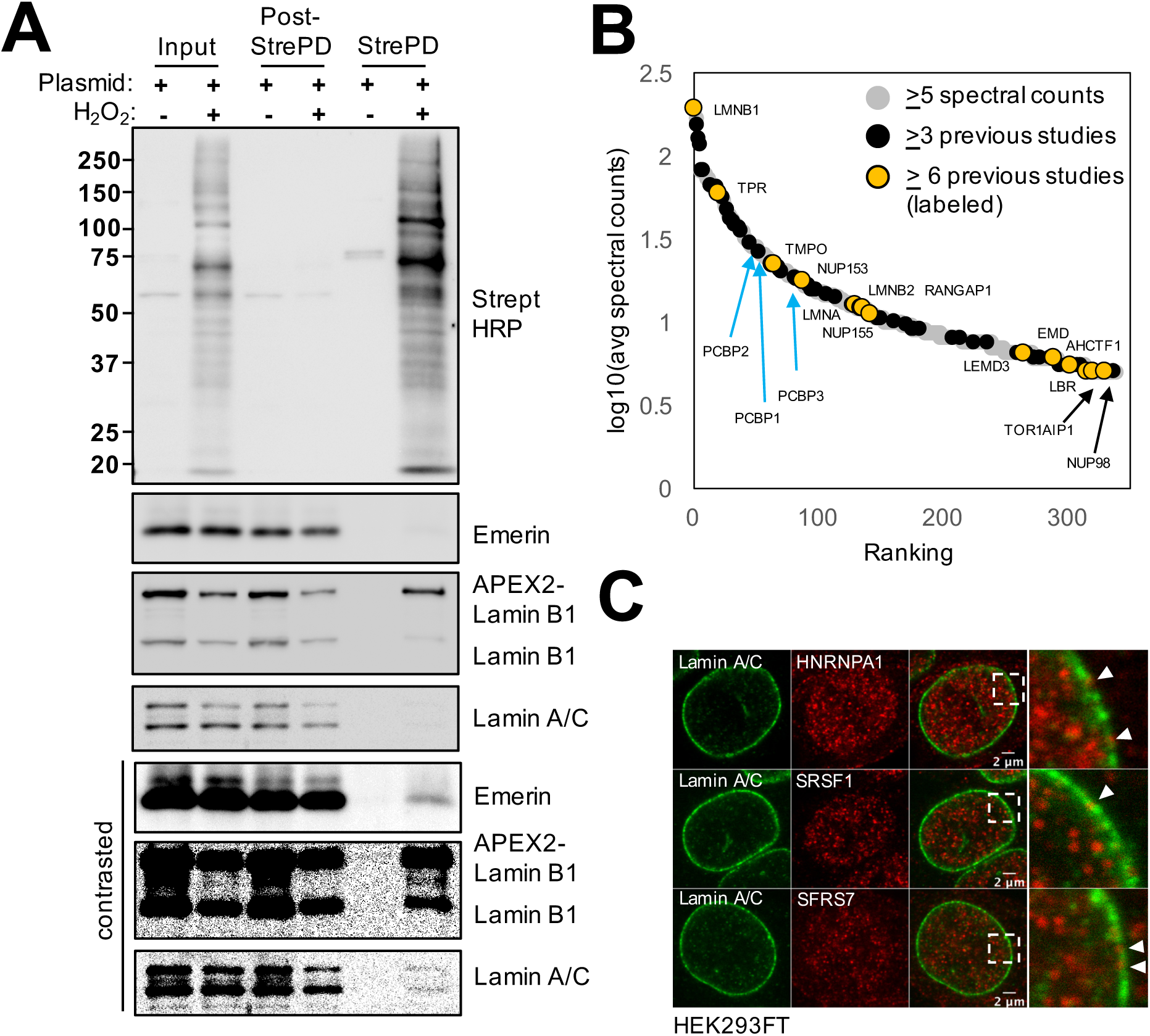
APEX2-lamin-B1 labeling identifies the proteome of the NL in HEK293FT cells. (A) Western blot for the APEX2 reaction with or without hydrogen peroxide and the streptavidin pull-down procedure. Detection was performed for biotinylated proteins (Streptavidin HRP) and indicated nuclear lamina components (Emerin, lamin-B1 and lamin-A/C) using the input, post-Streptavidin pulldown (post-StrePD) and Streptavidin pulldown (StrePD) fractions. A high contrast version of the Emerin, lamin-B1 and lamin- A/C blots is presented below the original images to better visualize the pull-down of NL components and the control experiment. (B) A graphical representation of the mass-spectrometry candidates plotted according to abundance rank and log10 of the average number of spectral counts. Grey circles represent proteins with an average of 5 or more spectral counts. Black circles represent proteins identified in 3 or more previously published NL proteome studies and the yellow circles with associated labels represent those seen in 6 or more studies. Blue arrows point to the ranking of C-rich element binding protein PCBP1-3. (C) Localization of select NL candidates (HNRNPA1, SFSR1 and SFRS7) from our NL proteome study. Inset represents a zoomed in view and arrowheads point to puncta that are juxtaposed to the NL.

A GO-term analysis of our NL proteome dataset revealed a strong enrichment for proteins involved in RNA splicing/stability, nuclear protein localization and DNA replication (Supplementary table 2). We identified the five components (IGF2BP1, HNRNPU, SYNCRIP, YBX1, and DHX9) of the CRD-mediated mRNA stabilization complex involved in β-catenin mediated *C-MYC* RNA stability (Noubissi et al., 2006; Weidensdorfer et al., 2009); and seven members of the T-chaperonin complex (CCT, TCP1, CCT2, CCT3, CCT4. CCT6A, CCT7, and CCT8), which contributes to protein folding and the localization of proteins in nuclear sub-regions such as telomeres and Cajal bodies (Freund et al., 2014; Gestaut et al., 2019; Wrighton, 2015). We also identify proteins involved in the initiation of DNA replication (MCM2/4/6/7, RFC1/3 and RPA1), a process that is known to be negatively affected by lamin mutants (Moir et al., 2000).

Since we observed an enrichment for a C-rich sequence (CCCWCCC) in the 3’ UTR in our APEX NL-RIP experiments above (Figure 2H), we anticipated that our NL proteome experiments would identify a protein or a complex of proteins that could bind polyC (“poly(rC)”) sequences. Indeed, our NL-proteome contains three known poly(rC) binding proteins, Poly(rC) Binding Protein-1 (PCBP1), PCBP2 and PCBP3, which had abundances greater than other known NL proteins such LMNA and EMD (Supplementary table 2, Figure 3B, blue arrows). The NL proteome identified by APEX suggests a role for the NL in RNA regulation, and also identifies proteins at the NL that are involved in protein localization and DNA replication.

### APEX2-lamin-B1-based labeling identifies both stable and variable lamin-associated domains (LADs) in G1, S, and G2 cells

The protein-rich NL interacts with specific DNA regions known as lamina-associated domains (LADs) (Guelen et al., 2008). To see if APEX2 could be used to map LADs by streptavidin pull downs of DNA-protein complexes (hereafter referred to as APEX-ID), we performed the reaction in K562 cells, which is a readily transfectable cell line previously used for both DamID and TSAseq mapping of LADs (Chen et al., 2018b). We found that the K562 line contained LADs mapped by APEX-ID which were similar to DamID and TSAseq (both based on DNA labeling) (Figure 4A), with Pearson coefficients of 0.84 and 0.81 (Figure 4B), respectively. The distribution of LADs, which were defined by Hidden Markov Modeling (see below), across chromosomes is similar between all methods, and is most similar between the related APEX-ID and TSAseq methods (Figure S4A). Gene expression in APEX-ID defined LADs was much lower than those outside of LADs, as expected (Figure S4B). We conclude that APEX-ID reliably maps LADs in cultured cells.

**Figure 4:**
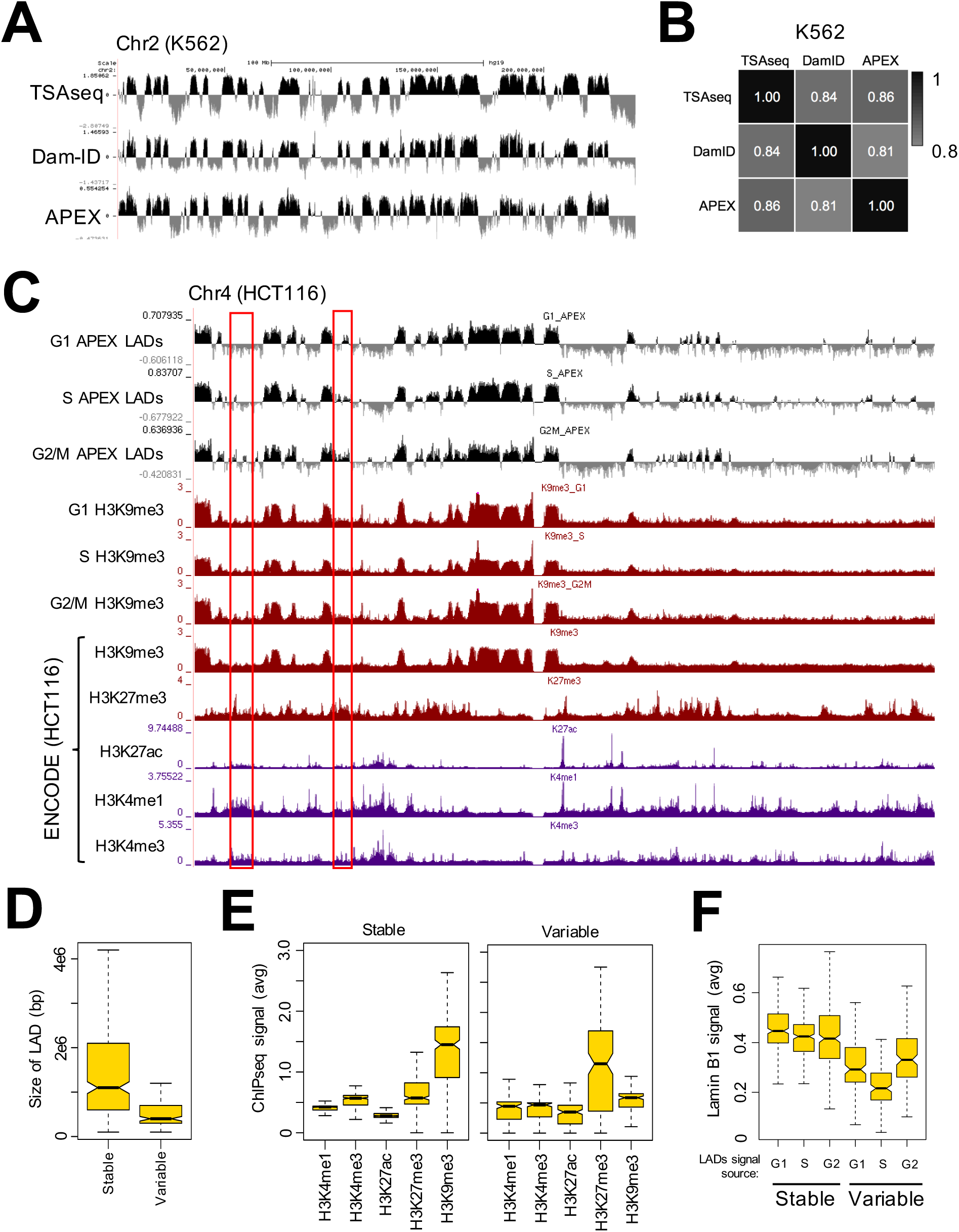
APEX-ID identifies lamina-associated domains (LADs) (A) UCSC Genome browser tracks for DamID, TSAseq and APEX-ID LADs data on chromosome 2 (hg19) for the K562 cell line. (B) Pearson correlation heatmap for K562 LADs data obtained by DamID, TSAseq and APEX-ID. The data were binned into 100kb genomic windows and averaged. (C) UCSC Genome browser tracks for HCT116 cell cycle APEX-ID LADs and H3K9me3 ChIP-seq data obtained in this study, and HCT116 ENCODE data for H3K4me1, H3K4me3, H3K9me3, H3K27ac and H3K27me3. (D) Boxplot showing the size range of stable and variable LADs. (E) Boxplot showing the average signal for ENCODE H3K4me1, H3K4me3, H3K9me3, H3K27ac and H3K27me3 ChIP-seq data in stable (left) and variable (right) LADs. (F) Boxplot showing the log2(StrePD/input) signal from each cell cycle stage (“LADs signal source”) in stable and variable LADs groups. The notches in these boxplots represent the 95% confidence interval around the median.

LAD mapping is typically done in asynchronous cell populations and it is not clear exactly how similar LADs are in different stages of the cell cycle. To determine this, we performed APEX-ID in HCT116 cells, a cell line often used in studies of mitosis. Fixed cells are FACS sorted for G1, S, and G2/M populations and then subjected to the APEX procedure. Since the NL disassembles in M phase, LAD maps for the G2/M population correspond to the G2 phase. We normally observed ∼40-60% APEX2-lamin-B1 transfection efficiency and sorted for ∼1 million cells in each cell cycle stage. LAD maps of each cell cycle stage reveal no large changes in profiles across G1, S, and G2 phases (Figure 4C). Next, we carried out chromatin immunoprecipitation sequencing (ChIP-seq) for H3K9me3, which is abundant in heterochromatin/LADs. The results show that the pattern for H3K9me3 remains largely the same throughout the cell cycle, and these data correlated well with cell cycle LADs data (Figure 4C, S4C). Further, a cross-referencing of our LADs and H3K9me3 datasets with the H3K9me3, H3K27me3, H3K27ac, H3K4me1 and H3K4me3 ENCODE datasets for HCT116 cells indicate that our LADs/H3K9me3 datasets are reliable with positive correlation with repressive chromatin (H3K9me3, H3K27me3) and negative correlation with active chromatin marks (H3K4me1, H3K4me3 and H3K27ac; Figure S4C).

Although LADs patterns are grossly similarly across G1, S, and G2, we noticed some interesting variability (Figure 4C, red boxes). To further analyze this variability, we first defined LADs using a three-state Hidden Markov Model (HMM). The three-state model is used to distinguish between (1) strong LAD signals, (2) an intermediate LAD signal that is characterized by a mixture of both enrichment and lack of enrichment or a weak signal, and (3) a signal that is clearly not LADs. The presence of intermediate-type signals suggests that APEX-ID, like TSAseq, produces less binary data than DamID (Chen et al., 2018b). Intersecting HMM LADs between experimental replicates result in an average of ∼85% overlapping LADs calls for each cell cycle stage, and yielded 378 G1-, 303 S- and 389 G2-phase LADs (Figure S4D). The average number of HMM LAD calls in S-phase is slightly reduced across virtually all chromosomes (Figure S4E). When we compare LADs in each cell cycle, we find that >80% of LADs are shared among all cell cycle stages (Figure 4D). These stable LADs have an average size of ∼1.5 megabases and are strongly enriched for the heterochromatin marker H3K9me3 (Figure 4D, 4E). LADs that are different between any cell cycle stage, which we call variable cell cycle LADs, tend to be smaller in size (average ∼500kb, Figure 4D), and are instead strongly enriched for H3K27me3 (ENCODE data, Figure 4E). Finally, we find that the average lamin-B1 log2(StrePD/input) signal was reduced in variable LADs across the cell cycle and was most pronounced during S-phase (Figure 4F). Thus, the APEX-ID reveals that LAD regions enriched for H3K9me3 are stable structures throughout the major stages of the cell cycle and that LADs variability exists in regions enriched primarily for the heterochromatin marker H3K27me3.

## Discussion

In an effort to more comprehensively examine the NL and to gain insights into its functions, we employed the APEX2 enzyme fused to lamin-B1 to map the proteins, RNA and DNA associated with the NL. We find that the APEX2 enzyme is robust and can be used in both live cells and PFA-fixed cells, making it amenable to numerous applications. The robust APEX-mediated biotinylation under the fixation condition is particularly advantageous since it greatly reduces protein diffusion and thus restrict biotinylation to the NL. This feature, in conjunction with a fast (∼20 seconds) APEX reaction, should allow the study of how NL-associated RNA and chromatin may change in different cell cycle stages.

In this study, we used APEX2 to identify RNAs associated with the NL. Our findings suggest a potential role for the NL in facilitating removal of introns, or alternatively the retention of some intron-containing transcripts for degradation (Boutz et al., 2015; Lareau et al., 2007). The presence of RNAs with introns at the NL, and the reduction of such introns in RNAs in the cytosol were previously described in the APEX-RNA-seq experiments based on labeling of RNA from the Ting lab and is an avenue of exploration in lamin biology (Fazal et al., 2019). Interestingly, we found that the NL associates with a small subset of highly expressed mRNAs. These mRNAs contained long 3’ UTRs suggesting a role for the NL in the regulation of mRNAs containing this feature. This is supported by a finding that the deregulated genes in lamin-null (TKO) mESCs also contain long 3’ UTRs. Subsequent analysis of these 3’ UTRs identified a C-rich motif (CCCWCCC) as a potential regulatory element in both the NL-enriched RNA and in RNAs deregulated in the TKO mESCs. C-rich motifs were previously found to be involved in alpha-globin and mu-opioid receptor mRNA stability (Hwang et al., 2017; Kong and Liebhaber, 2007). Further, the length of the 3’ UTR can affect RNA stability and localization by providing a platform for other regulatory elements, such as microRNAs or through the use of alternative polyadenylation sites (APA) (Mayr, 2016). The length of the 3’ UTR, and in particular the use of APA sites, is implicated in a number of pathologies including cancer and cell senescence, and in cellular stress response (Chang et al., 2015; Chen et al., 2018a; Mayr and Bartel, 2009). Senescence of cultured primary cells, for example, is a phenomenon linked to NL biology and to the reduction of lamin-B1 (Freund et al., 2014; Shimi et al., 2011).

We show that the NL proteome mapped by APEX2-lamin-B1 enriches for both known NL components and proteins involved in RNA processing, protein localization and DNA replication. Especially relevant to this study are the poly(rC) binding proteins. Poly(rC) proteins have been reported to affect the stability of RNAs, and they also appear to more broadly affect the transcriptional landscape of cancer cells (Behm-Ansmant et al., 2007; Perron et al., 2018). These proteins could regulate the stability of mRNAs containing 3’UTR C-rich motifs at the NL, thus explaining some of the genes that are deregulated in the TKO mESCs (Zheng et al., 2018). It would be interesting to further establish the role of NL in mRNA stability via interacting mRNA and poly(rC) binding proteins (Choi et al., 2009).

Our analyses of the datasets presented in this study, for which there is precedent, also highlights both methodological and biological variation, both of which are to be expected. For example, our data is largely but not completely concordant with that mapped using lamin-A/C and the BAR method (Bar et al., 2018). Further, the RNAs identified in our study do not strongly overlap (∼1.5%) with those identified by the Ting group using APEX2-lamin-A/C via pulling down RNA directly (Fazal et al., 2019). We believe that some of this variability could be attributed to the observation that lamin-A/C and lamin-B1 form distinct meshworks in human and mouse tissue culture cells (Schermelleh et al., 2008; Shimi et al., 2008). The previous studies used lamin-A/C while here, we used lamin-B1. The variability related to the RIP experiment can also be attributed to the streptavidin pull down of labeled RNA (Fazal et al., 2019) versus pull down of RNA via its associated proteins. It might be expected that RNAs stably associated with the NL are more readily labeled than RNAs *en route* to the cytosol that are often shielded by proteins. These latter RNAs would be enriched in our method but not by the method reported by the Ting group (Fazal et al., 2019). Alternatively, variability between studies might also reflect true differences among different cell types and would be of future interest considering the tissue specificity of the many laminopathies (Hatch and Hetzer, 2014; Schreiber and Kennedy, 2013).

We show that APEX2 can be used to reliably map LADs during dynamic processes such as the cell cycle. Our results indicate that most LADs are quite stable during the cell cycle. These stable LADs were generally large in size (∼1.5 megabases), and most strongly enriched for the heterochromatin mark H3K9me3. On the other hand, LADs that were variable between cell cycle stages were smaller (0.5 megabases) and were primarily enriched for the heterochromatin mark H3K27me3. We also found that variable LADs showed a reduced lamin-B1 signal, and that this was especially pronounced during S-phase of the cell cycle. The explanation for the latter is unclear but may be related to DNA replication. Our previous modeling studies of histone lamin landscapes (HiLands) in mESCs identified two types of LADs, HiLands-B and P (Zheng et al., 2015), that were largely concordant with the facultative and constitutive LADs, respectively (Meuleman et al., 2013; Zheng et al., 2015). Interestingly, HiLands-B LADs have similar features to the cell cycle variable LADs we identified here. HiLands-B LADs are short and exhibit weak lamin-B1 signal, and enrich for H3K27me3. On the other hand, the HiLands-P LADs are similar to the cell cycle constitutive LADs we describe here, and are larger in size and H3K9me3 rich. It will be important to further investigate the similarities between cell cycle variable and stable LADs we found here and HiLands-B and -P LADs defined in mESCs. Interestingly, we observed that lamin deletion caused HiLands-B detachment from the NL and HiLands-P decompaction at the NL in cell cycle asynchronous mESCs (Zheng et al., 2018). Based upon these considerations, it would be important to understand if the differential changes of HiLands-B and -P LADs are related to cell cycle states in lamin-TKO mESCs (Zheng et al., 2018).

The mapping techniques described in this study enabled by the APEX method are ideal for applications requiring genomic, RNA and/or protein interactions located in a defined cellular structure such as the NL. In the future it will be of interest to carry out temporal studies of changes in protein composition, RNA, NL, and LADs in responses to a cellular stress such as a mechanical perturbation or chemical response are particularly good candidate systems. Further, the APEX2-linked system is an ideal platform for situations where a discriminating antibody is unavailable, such as when dealing with genetic variants and closely related isoforms. In this study, we present a flexible and temporally capable method of characterizing the NL structure that we believe will be a valuable tool.

## Acknowledgements

We would like to thank members of the Zheng and Goldman labs, and Matthew Sieber for advice and discussions. We also thank Allison Pinder, Frederick Tan and Xiaobin Zheng for their help with sequencing and data analysis. This study was funded by NIH NIGMS (GM106023) to Drs. Robert Goldman and Yixian Zheng, NIH NIGMS (GM110151) to Yixian Zheng, and NIH NIGMS (8 P41 GM103533) to John R. Yates III.

---

**Figure S1.**
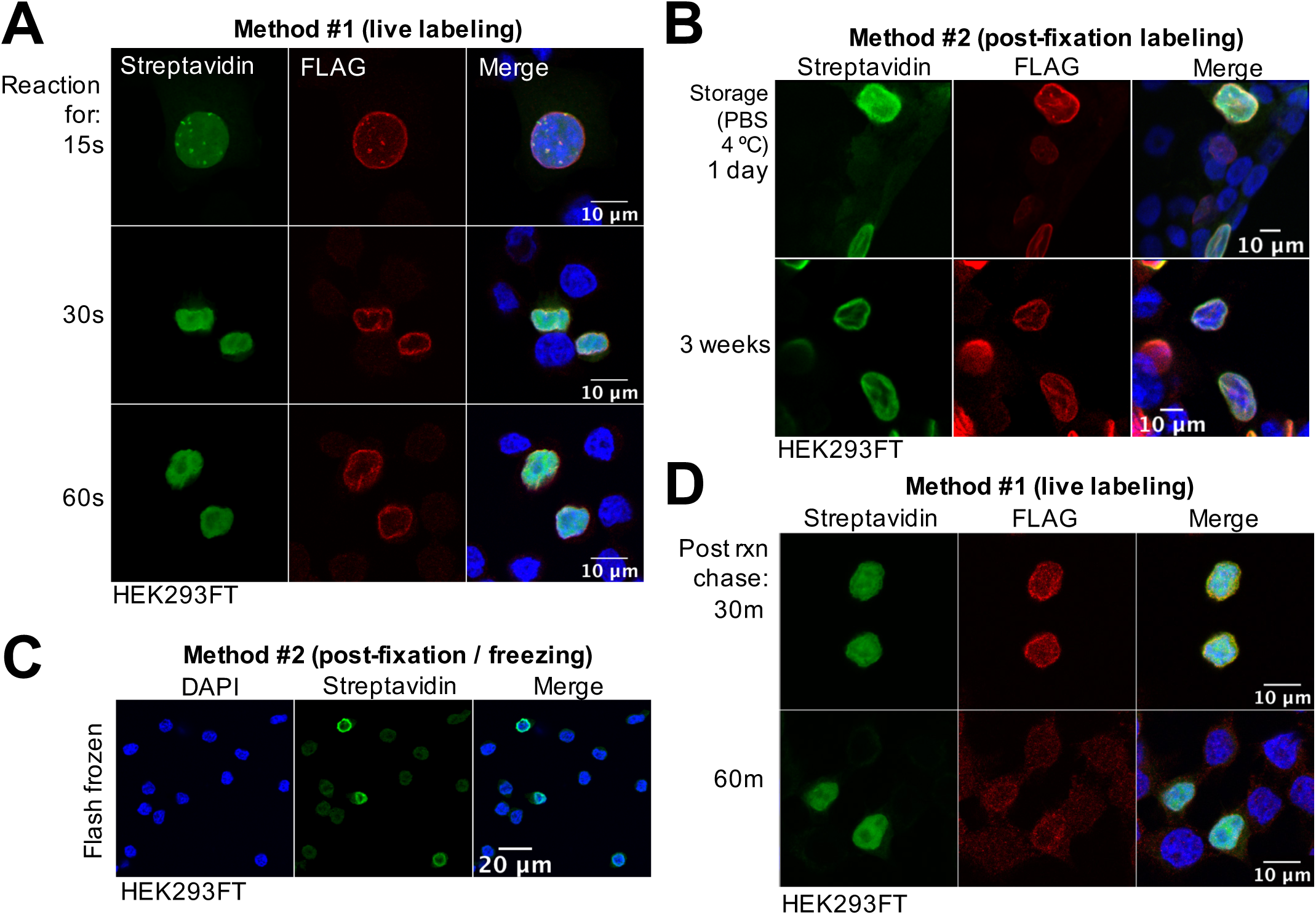
(related to Figure 1): The APEX2 reaction in APEX2-lamin-B1 cells is quick, durable and labels material that remains in the nucleus. The APEX2 experiment performed for (A) different lengths of time in live HEK293FT cells, (B) with HEK293FT cells that were fixed and stored in PBS for one day or for three weeks, (C) after PFA-fixation and snap freezing and (D) as a pulse-chase experiment in live HEK293FT cells that were then fixed with PFA for fluorescence staining. All samples were stained with DAPI (DNA), Streptavidin (biotinylated material) and FLAG M2 (APEX2-lamin-B1).

**Figure S2.**
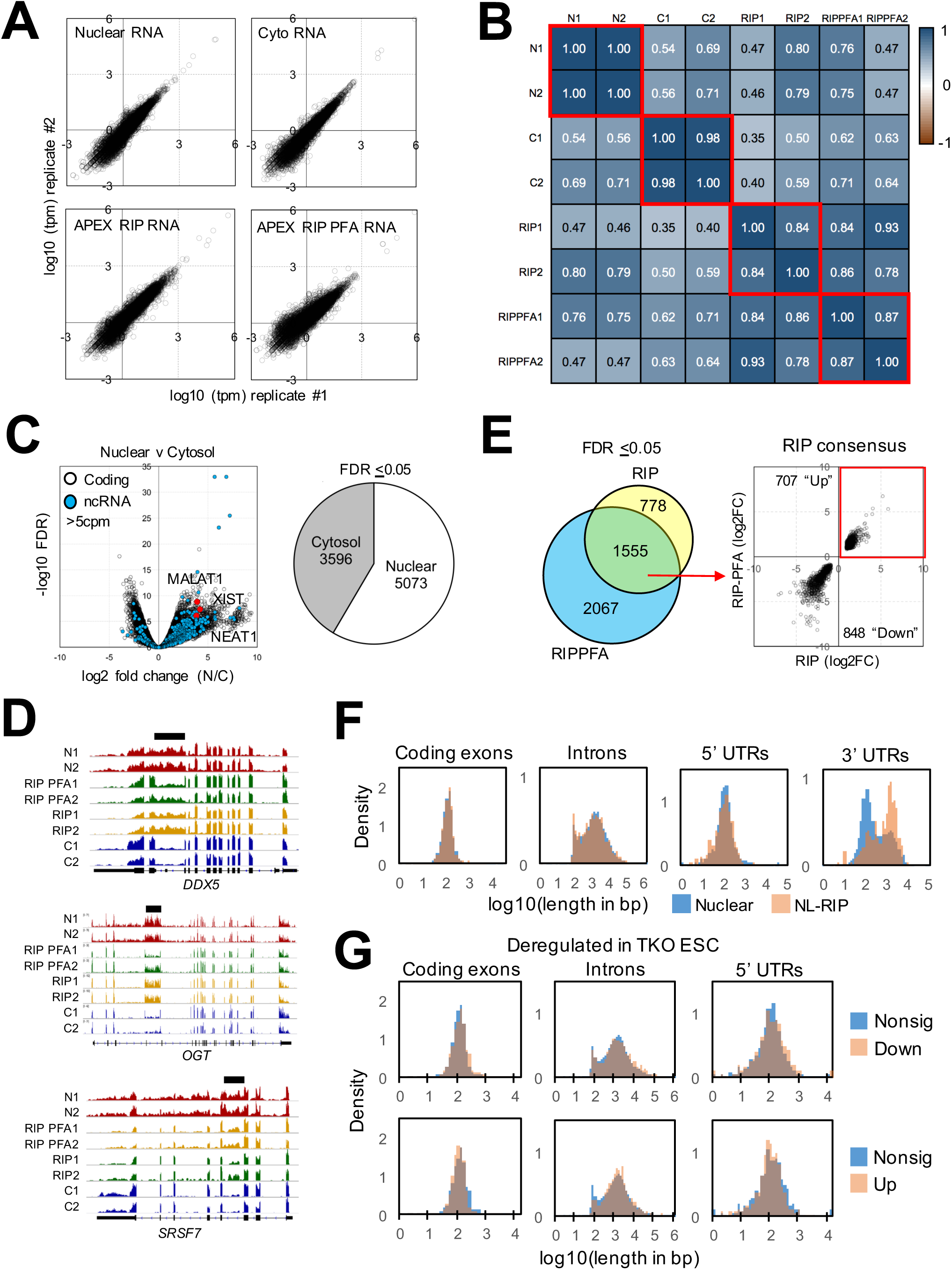
(related to Figure 2): APEX RIP biological replicates and enrichment analysis. (A) Plots of replicate HEK293FT RNA-seq datasets. Transcripts per kilobase per million (TPM) were transformed as log10 value. (B) A heatmap matrix representing the Pearson correlation of the TPM between each dataset. Red boxes highlight replicate experiments. (C) Left panel is a Volcano plot showing the differential nuclear and cytosolic enrichment of RNAs. Black circles represent coding RNAs, blue circles represent noncoding RNA species and red circles indicate RNAs (MALAT, XIST, NEAT1) with known enrichment in the nucleus. Right panel is a pie chart illustrating the number of RNA differentially enriched in the nucleus and cytosol using an FDR of less than or equal to 0.05. (D) IGV browser view showing examples of RNA with known retained introns in our RNA-seq datasets. (E) Left panel is a Venn diagram showing the overlap (RIP consensus) between APEX-RIP (“RIP”) and APEX-PFA RIP (“RIPPFA”) RNA-seq datasets. Differential enrichment was determined against the nuclear fraction. Right panel is a plot of the log fold change directionality for the consensus datasets. We focused on the 707 enriched (“up”) RNAs. (F) Density histograms comparing the distribution of sizes for coding exons, introns, 5’ UTRs and 3’ UTRs between the NL-RIP and the nuclear enriched fraction. (G) Density histograms comparing the distribution of sizes for coding exons, introns and 5’ UTRs of deregulated genes in the lamin-null mESC RNAseq dataset.

**Figure S3.**
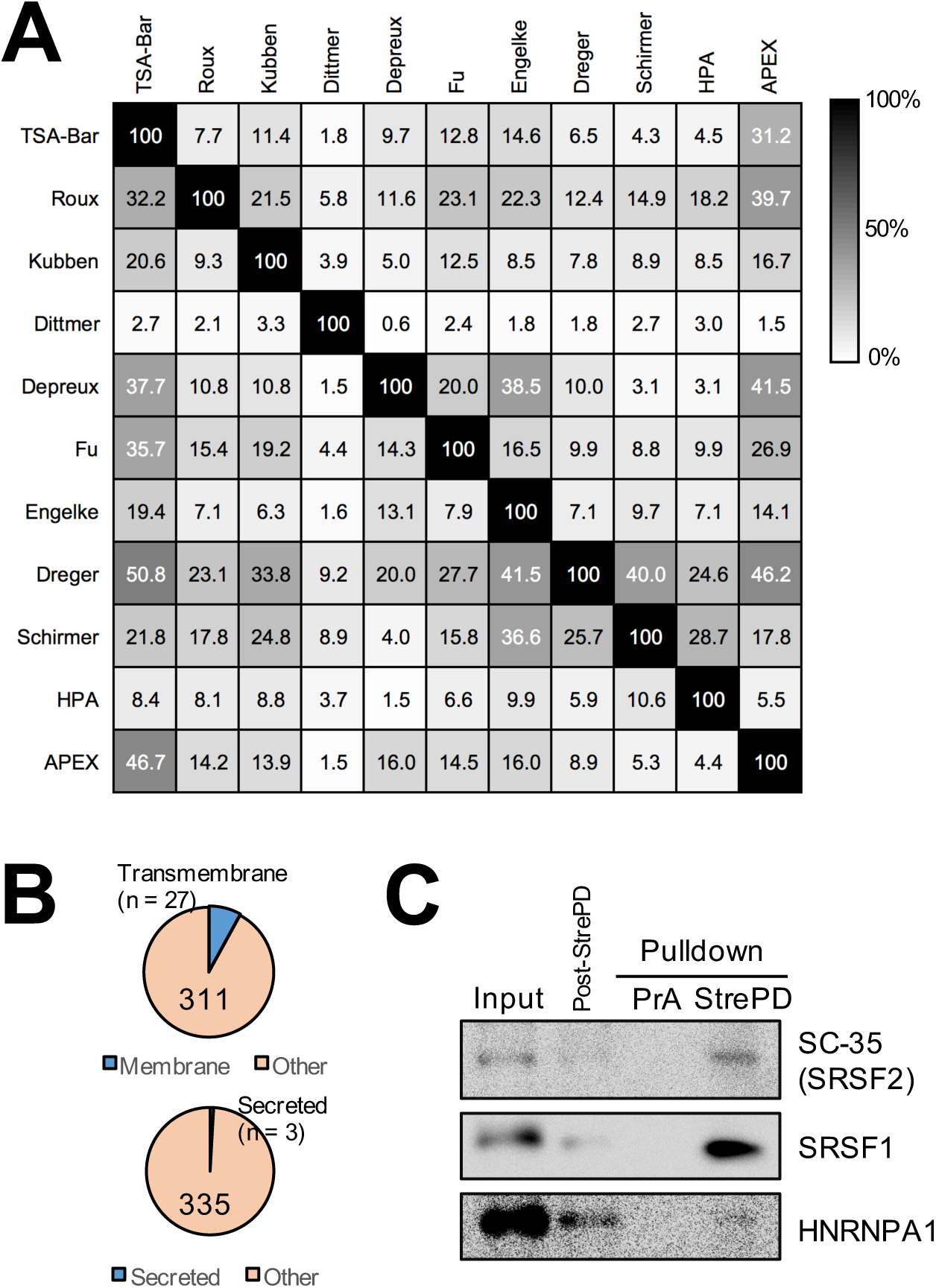
(Related to Figure 3): Comparison of our NL proteome with previous studies and visualization of select mass spectrometry hits. (A) A matrix representing the percentage overlap between the NL proteomes identified in previous studies. HPA represents nuclear membrane localized proteins identified by the Human Protein Atlas Project. Due to differences in the number of identified proteins from each study, the table is only read from left to right. (B) Pie charts of our NL proteome showing the number of transmembrane (top) and secretory (bottom) proteins as defined by the Human Protein Atlas Project. (C) Western blot validation of mRNA splicing related proteins (SRSF1, SRSF2, and HNRNPA1) identified in our NL proteome. PrA is a Protein-A dynabead control.

**Figure S4.**
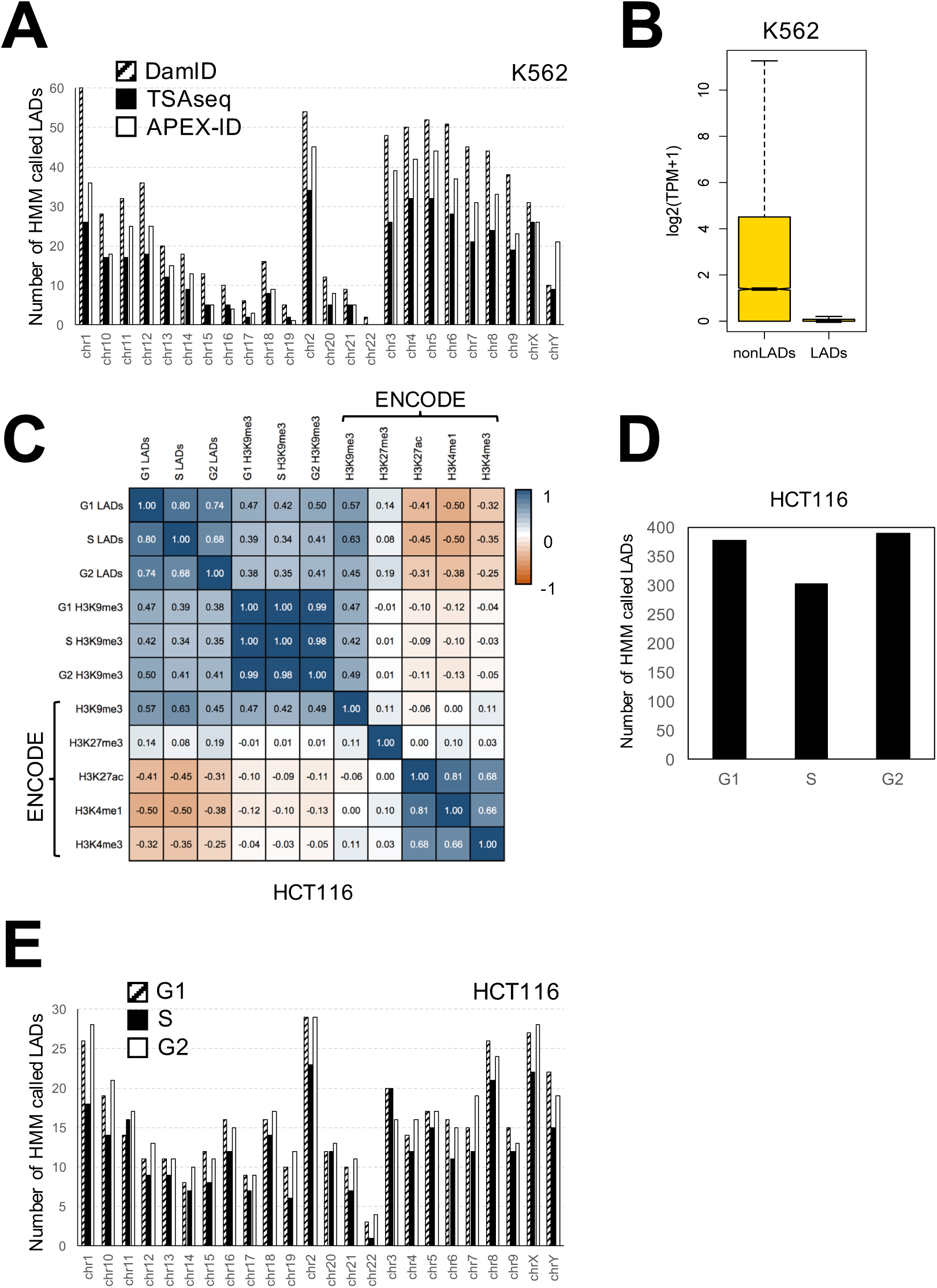
(related to Figure 4): APEX-ID LADs are correlated with features of heterochromatin and show minor variation throughout the cell cycle. (A) A graphical representation of the chromosomal distribution of K562 HMM-defined LADs from DamID, TSAseq and APEX-ID. (B) Boxplot showing the range of gene expression levels in and outside of K562 APEX-ID LADs. The notches represent the 95% confidence interval around the median. (C) Pearson correlation heatmap comparing our cell cycle HCT116 APEX-ID LADs and H3K9me3 ChIP-seq datasets and HCT116 datasets obtained from the ENCODE project. (D) Chart representation of the average number of LADs called by the Hidden Markov Model in each cell cycle stage from HCT116 cells. (E) The distribution of LADs calls across each chromosome during the HCT116 cell cycle. Notches represent the 95% confidence interval around the median.

